# Transcripts from multicopy gene families localizing to mouse Y long arm encode piRNAs and proteins

**DOI:** 10.1101/407197

**Authors:** Rachel A Jesudasan, Kankadeb Mishra, Pranatharthi Annapurna, Anurag Chaturvedi, Nissankararao M Praveena, Jomini L Alex, Sivarajan Karunanithi, Hemakumar M Reddy

## Abstract

Heterochromatic long arm of mouse Y chromosome harbors the multicopy species-specific sequences *Ssty, Sly, Asty,* and *Orly* that are transcribed in testis. Of these *Ssty* and *Sly* genes encode proteins – yet all the copies of these RNAs are not translated. Using bioinformatic approaches, small RNA northern blots and electrophoretic mobility shift assays, we demonstrate here that these multicopy gene families from mouse Y-long arm generate piRNAs predominantly in testis. Thus, we identified a piRNA cluster on mouse Y chromosome and also unraveled the dual role of Y-chromosome-encoded transcripts to act as primary transcripts of piRNAs in addition to their role as protein-coding RNAs.

**HIGHLIGHTS:**

1. First report of a cluster of piRNAs on a mammalian Y chromosome
2. Report of primary transcripts of piRNAs
3. These piRNAs putatively regulate autosomal genes expressed in mouse testis Ssty and Sly genes code for proteins as well as generate piRNAs

## INTRODUCTION

The role of mammalian Y chromosome in testis-determination, spermatogenesis and male fertility(1-3) is well established. The Y-chromosomes in general contain few genes and are replete with repeats(4-6). The entire long arm of the Mouse Y-chromosome is heterochromatic and harbors a unique repertoire of coding and non-coding, sex and species-specific multi-copy genes that are expressed testis-specifically(7, 8). The testis-specific *Ssty* (spermiogenesis-specific transcript on the Y), *Sly* (Sycp3-like, Y-linked), *Asty* (amplified spermatogenic transcripts Y encoded) and *Orly* (oppositely-transcribed, rearranged locus on the Y), are X-Y homologous multicopy genes present on the heterochromatic long arm of the mouse Y chromosome (8-11).

There are more than 300 copies of *Ssty* with two distinct intronless subfamilies, *Ssty1* and *Ssty2.* They are ∼84% identical at the nucleotide level on mouse Yq(8). *Sly* on the other hand has protein coding and noncoding copies (9, 11). Several peptides have been predicted from long noncoding RNAs(12). SSTY1 and SLY proteins are expressed in mouse spermatids(9). SLY is thought to be involved in multiple processes in spermiogenesis as it interacts with an acrosomal protein, a histone acetyl transferase and a microtubule-associated protein. The *Asty* and *Orly* transcripts do not appear to have coding potential. Large numbers of Orly isoforms are perhaps transcribed at low levels(7).

Mice with deletions of long arm of Y chromosome show, sperm morphological and motility related abnormalities concomitant with the extent of deletion(9). These mice also exhibit either subfertility/sterility that is dependent on the extent of deletion. Deletions of long arm of the mouse Y chromosome, also lead to upregulation of a number of X- and Y-transcripts in spermatids(13). The proteins encoded by the *Ssty* and *Sly* gene(8) families are envisaged to be responsible for the phenotypes observed in mice with deletions of Yq(3, 14).

Although some copies of the *Ssty1, Ssty2* and *Sly* code for proteins, others are transcribed but not translated(7-10). Functional significance of these noncoding transcripts is not known. Earlier we demonstrated that a stretch of 67 nucleotide sequence transcribed from the distal block of heterochromatin from human Y long arm functions as the 5’ UTR of an autosomal gene in testis(15). Therefore, in an attempt to find functions for the mouse Y-transcripts, we queried the UTR database. *In silico* analysis identified short stretches of sequence homologies between the Y-transcripts and UTRs of many genes localizing to autosomes. Small RNA northern blotting and gelshift assays using oligonucleotide probes corresponding to the homologous sequences identified piRNAs. Thus, in the present study, we provide experimental evidence to support generation of piRNAs from the multicopy gene families localizing to the mouse Y chromosome for the first time.

## RESULTS

### Transcripts from multicopy gene families on mouse Yq generate piRNAs in testis

BLAST analysis of the transcripts derived from multicopy genes on mouse Y chromosome such as *Sly, Ssty1, Ssty2, Asty,* and *Orly* against the UTR database identified 123 short stretches of homology (17-30 nucleotides), in 3’ UTRs and 5’ UTRs of many X-chromosomal and autosomal genes across a number of species from viruses to humans, in both +/+ and +/-orientations (Table 1). Coding sequences from mouse transcriptome were extracted from <u>Ensembl</u>. BLAST analysis of the Y-transcripts against the coding sequences (CDS) did not identify any homology except for the parent transcripts *Sly* and *Ssty2*.

**Table 1.** BLAST analysis of Y transcripts against <u>UTRdb</u>

| Y transcripts | Number of motifs in UTRs |  | Number of motifs in either Orientation |  | Chromosome localization of target genes (*Mus musculus*) |
| --- | --- | --- | --- | --- | --- |
|  | 3' | 5' | +/+ | +/- |  |
| *Sly* | 11 | 3 | 10 | 4 | 7 |
| *Ssty1* | 18 | 11 | 21 | 8 | 1, 7, 12 & 16 |
| *Ssty2* | 15 | 2 | 10 | 7 | 11 |
| *Asty* | 11 | 3 | 9 | 5 | 2, 3, 5, 9 & 17 |
| *Asty-sv* | 33 | 4 | 23 | 14 | 4, 9, 11, 12 & 13 |
| *Orly* | 11 | 1 | 5 | 7 | 8 |
| Total | 99 | 24 | 78 | 45 | - |

*BLAST analysis of Y-transcripts against the mouse UTRs reveals short stretches of identity in the 5' and 3'UTRs of genes localizing to different chromosomes in both +/+ and +/- orientations, indicating interactions between the Y-chromosomal and autosomal genes.*

Locked nucleic acid (LNA) oligonucleotide probes (11-23 nt) homologous to different UTRs that have matches to the Y-transcripts were used as probes on small RNA blots (20 μg in each lane). The probes were checked for homology to mouse X chromosome. Only the probes specific to the Y were chosen for northern blot analysis. Fifteen of the seventeen probes identified 26-32 nt long signals predominantly in testis, which suggested that these could be piRNAs based on their size (Fig. 1a-n and p); Orly.2 and Orly.4 did not identify testis-specific signals even with 30 μg of small RNA (Fig. 1 o and q). Orly.3 identified a faint signal in testis, on a 50 μg small RNA blot (Fig. 1p) - indicating low levels of transcription of these RNAs. The low level of expression of the *Orly* isoforms is perhaps the reason for barely detectable levels of these piRNAs. We observed piRNAs mainly in testis although two of the oligonucleotides identified signals corresponding to piRNAs in brain and kidney respectively (Fig. 1i and j). The localization of probes used for small RNA northern blotting is depicted on the corresponding Y-transcripts (Supplementary Fig. 1). A scrambled probe that was used as negative control did not give any signal in the range of small RNAs (∼26-32 nucleotides) in any of the tissues (Fig. 1r) indicating that the signals observed with the other probes are specific. U6 is a highly conserved, ubiquitously transcribed, low molecular weight small nuclear RNA. A probe corresponding to U6 RNA served as the loading control for all the blots(16).

**Figure 1.**
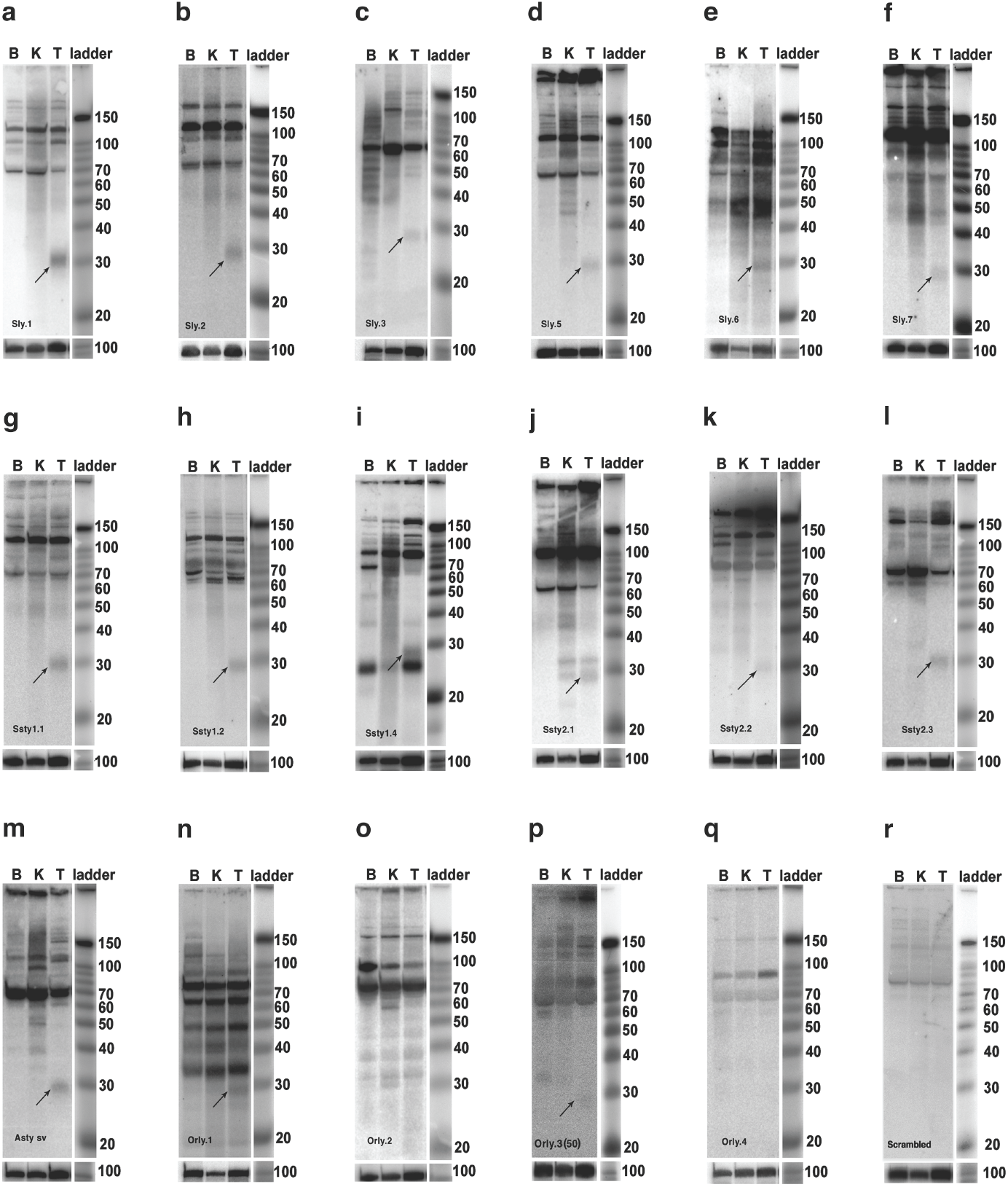
Small RNA northern blots showing signals corresponding to piRNAs. LNA oligonucleotides showing homology to Y-chromosomal transcripts and UTRs of different genes-were used as probes on northern blots containing 20 μg small RNA from brain (B), kidney (K) and testis (T). (a-f) Testis-specific signals of 29-30 nt observed with all Sly probes. (g-m) LNA probes from Ssty1, Ssty2 and Asty-sv show signals ranging from 26-32 nt. (i) In addition to the testis-specific signal at ∼29 nt, Ssty1.4 yielded signals of 26 nt in all tissues; (j) Ssty2.1 shows two bands of 29 and 31 nt in both kidney and testis; (o) Orly.2 shows a distinct band of ∼30 nt in kidney. (n, p) showed ∼30 nt testis-specific signals in Orly.1 and Orly.3 (50 μg blot). (o, q) Orly.2 and Orly.4 did not identify testis-specific signals of expected size (r) Scrambled probe used as negative control. U6 served as the loading control for all the blots.

In order to show that these small probes do not identify nonspecific signals, two of the Y-derived 11 nucleotide long probes Sly.1 and Ssty1.2 were used to examine expression in other tissues like male heart and spleen and female brain and ovary. The probes identified signals only in testis (Fig. 2a and b), which indicated the specificity of the probes.

**Figure 2.**
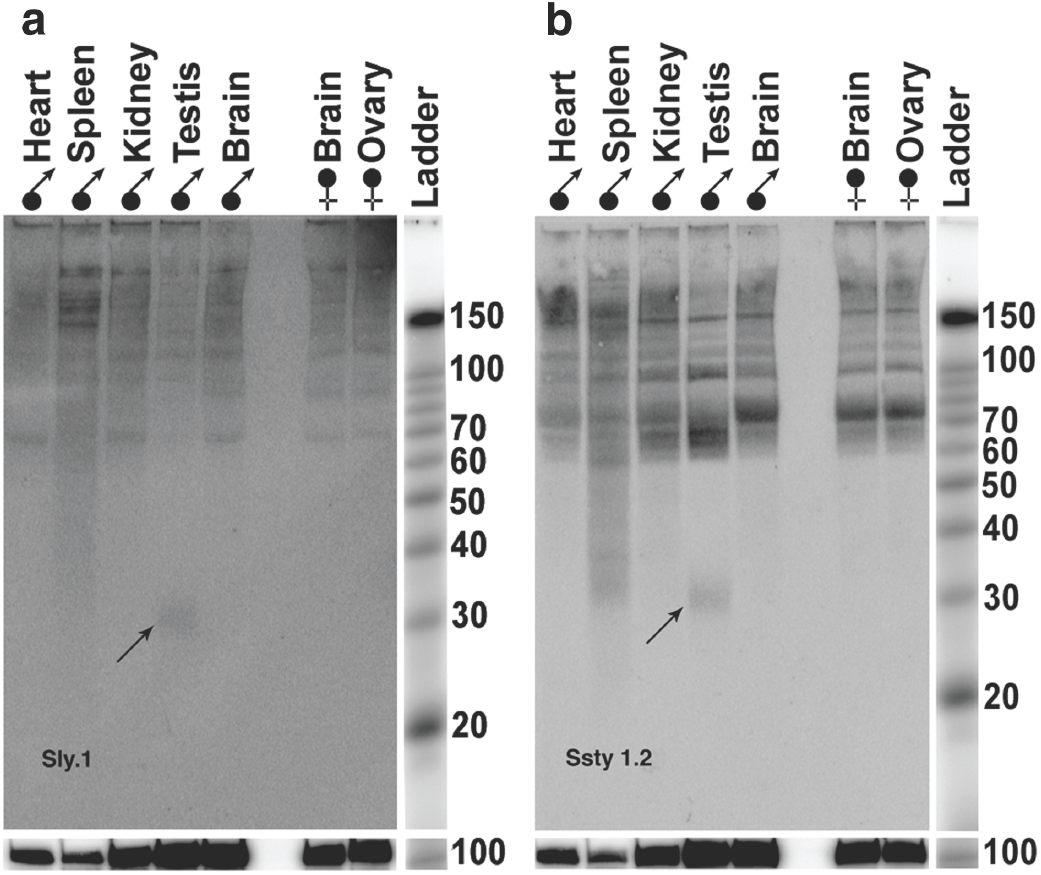
Multi-tissue small RNA northern blot. Northern blot including small RNA from male and female tissues (ovary and brain) showed ∼30 nt signals in testis only, with (a) Sly.1 and (b) Ssty1.2 probes. U6 was the loading control for both the probes.

piRNAs are a class of germline specific small RNAs that bind mammalian PIWI proteins(17). To further confirm that these RNA species are indeed piRNAs their ability to interact with MIWI protein was examined using Electrophoretic Mobility Shift Assay (EMSA). Recombinant MIWI protein (the mouse homologue of PIWI) was purified using pAAV-IRES-hrGFP vector and the specificity was confirmed by western blotting. MIWI antibody identified a single band of ∼98kDa in the testicular lysate and the purified fraction (Fig. 3a). RNA oligonucleotides were synthesized to the UTR homologous regions of *Sly* and *Ssty*. Binding of these to MIWI protein was checked using gelshift assays. Both showed shift in mobility upon incubation with purified MIWI protein (Fig. 3b and c). The mobility shift was competed out with either cold piR1, a known piRNA(17) or MIWI antibody. Similarly, cold *Ssty* and *Sly* oligonucleotides as well as the MIWI antibody competed out binding of piR1 to MIWI protein (Fig. 3b and c) confirming that *Ssty, Sly* derived oligonucleotides are piRNAs. To show that the small RNAs identified in the current study bind specifically to MIWI, an alternate antibody (argonaute 3 antibody) was used as a competitor in the gel shift assays. This antibody does not compete out the binding of MIWI protein to the oligonucleotides, indicating that these sequences correspond to MIWI-binding RNAs (piRNAs).

**Figure 3.**
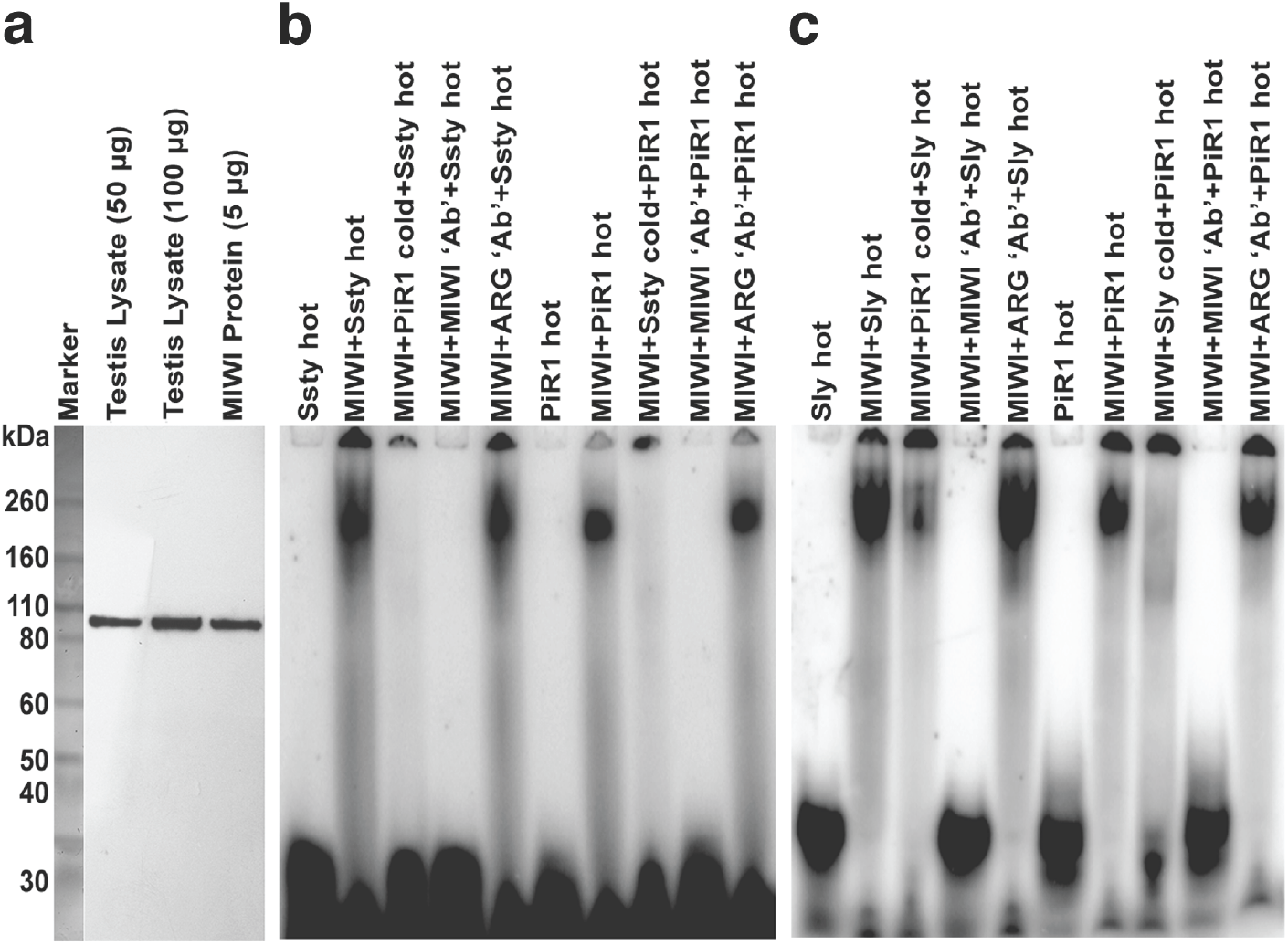
Putative piRNAs from mouse Y bind to MIWI protein. (a) Western blot analysis using mouse testis lysate and the purified recombinant MIWI protein with MIWI antibody showing ∼98kDa size protein in both. EMSA with MIWI protein and putative piRNA oligonucleotides from (b) Ssty and (c) Sly showed shifted bands similar in size to that of piR1; the gel-shifted bands were competed out on use of cold Sly/Ssty/piR1 oligonucleotides and MIWI antibody. A nonspecific antibody, Argonaute 3, did not compete out binding of the putative piRNAs to Sly/Ssty or piR1, indicating specificity of binding.

Computational analysis of the primary transcripts *Sly, Ssty1, Ssty2, Asty-sv* and *Orly* did not reveal any stable secondary structure, on using sliding-window technique with window size of 70 nt and window position increment of 1 nt (data not shown), using pknotsRG(18). Thus these transcripts do not have stable secondary structures that are characteristic of miRNA.

BLAST analysis using the Y transcripts (*Sly, Ssty* etc.) identified a total of 1166 small RNAs in the Sequence Read Archives at NCBI; <u>SRP001701</u> and <u>SRP000623</u> with 95% identity. The second dataset SRP000623 contained reads that have been annotated with respect to repeats. Of the 366 MIWI-MILI-associated reads identified in SRP000623 by the Y transcripts, 312 had homology to known repeats, and 54 did not. 11 of the 54 small RNAs were homologous to X-chromosome as well. Therefore, the 43 small RNAs that have homology only to the Y-transcripts could be derived from the Y chromosome.

We also observed that the genes showing sequence homology to the Y transcripts in their UTRs, showed expression in multiple tissues including testis and epididymis. These genes are expressed from different autosomes and X chromosome and have defined functions in the testis (Supplementary Table 1).

### Y-derived piRNAs are expressed differentially from the two strands of DNA

Oligonucleotide probes (Ssty1.4 and Orly.4), designed to both direct (sense, D) and complementary strands (antisense, C) yielded signals corresponding to the size of piRNAs with sense probes only, under identical conditions, revealing differential expression of piRNAs from the two strands (Fig. 4a and c). However, stringent washing and processing the blots in the sigmoid scale, elicited ∼30 nt long signals in testis with antisense probes also (Fig. 4b and d). Thus, both the sense and antisense strands of DNA produce piRNAs – albeit in unequal quantities.

**Figure 4.**
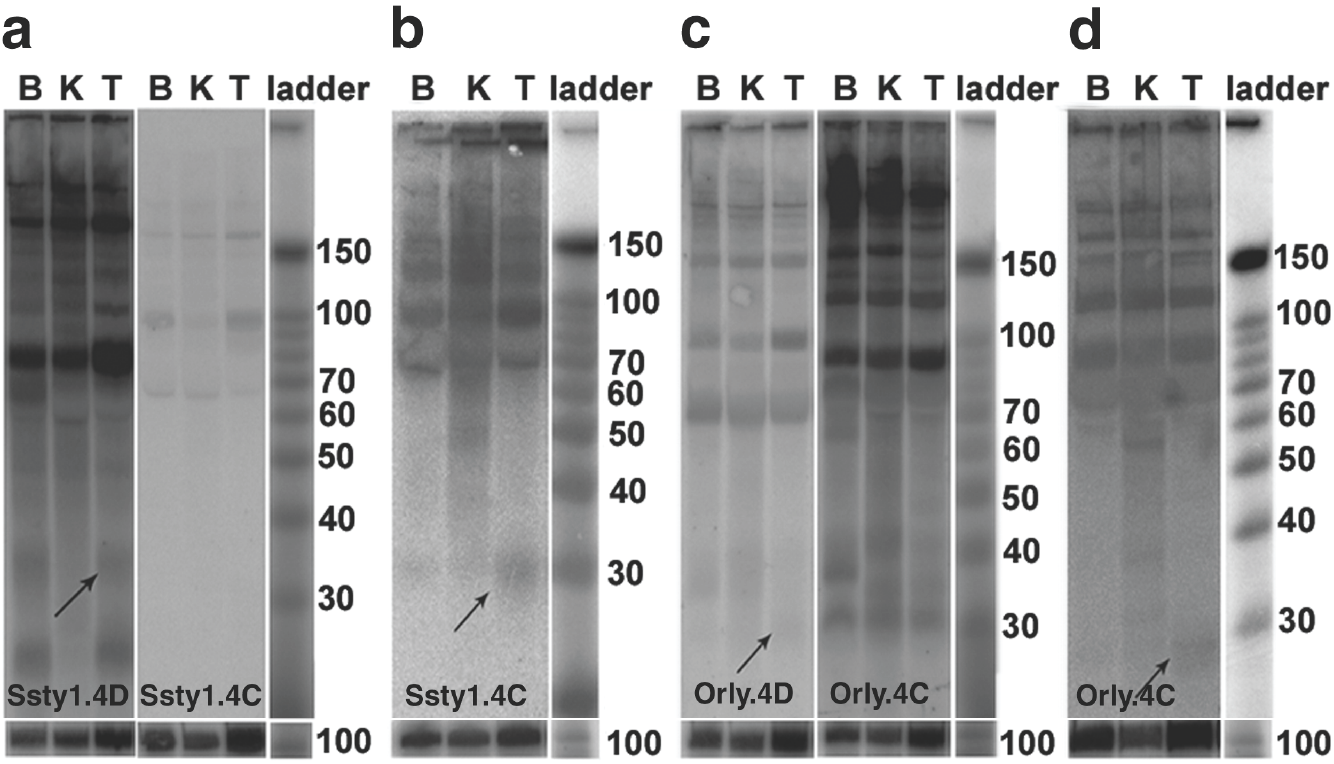
Small RNA northern blots showing differential expression of piRNAs from direct and complementary strands. Equal amounts of small RNAs from (B) brain, (K) kidney and (T) testis were electrophoretically separated, blotted, hybridized with probe noted on each blot, washed, developed and processed simultaneously. (a, c) LNA probes for predicted motifs from (D) direct and (C) complementary strands showed differential expression. (b, d) Blots hybridized with probes from complementary strands, revealed signals of approximately 30 nt in size, when washed under more stringent conditions and processed in the Sigmoid scale, indicating production of piRNAs from the complementary strand as well. U6 hybridization for each blot is given below the corresponding blot.

### Genes upregulated in Yq-deleted mice contain Y-derived piRNA motifs in their 3’ UTRs

Ellis and colleagues reported 23 genes that were upregulated in Yq-deletion mutant strains of mouse(13). These genes localized primarily to the X-chromosome with two of them showing autosomal localization. UTRs of nine of these upregulated genes contained short stretches of homology to the Y-transcripts (Table 2). Representative probes from two of the UTRs of upregulated genes, *H2al1* and *Xmr* (Sly-B, Ssty1-B), identified ∼30 nt testis-specific signals (Fig. 5A a and b). This indicated the presence of sequences homologous to piRNAs in these UTRs. A scrambled probe served as the negative control (Fig. 5Ac). Only two of the seven upregulated genes localizing to the X-chromosome (*H2al1, Xmr*) had homology to the Y-chromosome with coverage >80% and identity >80%; while five did not (Table 2).

**Table 2.** Genes that are upregulated in a Y-deletion mutant mouse (Ellis et al., 2005), with sequence homology to Y-transcripts in their UTRs.

| S. No. | Upregulated Gene | Chromosome Localization | Expression Pattern | Homology to Y – Chromosome |  | UTR homology |
| --- | --- | --- | --- | --- | --- | --- |
|  |  |  |  | %Coverage | %Identity |  |
| 1 | histone variant *H2al1* (AK005922) | X | T | >80 | >80 | 5' UTR<br>3' UTR |
| 2 | *Xlr*-related, meiosis regulated *Xmr* (NM_009529) | X | T | >90 | >90 | 3' UTR |
| 3 | Novel gene<br>Novel protein<br>Putative uncharacterized protein<br>Hypothetical LOC69357MCG1 13671 (AK005630) | X | T | -- | -- | 3' UTR |
| 4 | V-set and immunoglobulin domain-containing protein 1<br>Precursor (Cell surface A33 antigen)(Glycoprotein A34) (AK006251) | X | T/U | -- | -- | 3' UTR |
| 5 | Spermidine/spermine N(1)-acetyltransferase-like protein 1 | X | T | -- | -- | 3' UTR |
|  | (AK015086) |  |  |  |  |  |
| 6 | *TGFb*-induced factor homeobox 2-like, X-linked 1 (NM_153109) | X | T | -- | -- | 3' UTR |
| 7 | Tetraspanin-6 (*Tspan*-6)(Transmembrane 4 superfamily member 6) (NM_019656) | X | T/U | -- | -- | 3' UTR |
| 8 | Cystatin-9 Precursor (Testatin) (NM_009979) | 2 | T | -- | -- | 3' UTR |
| 9 | Glyoxylate reductase/hydroxypyruvate reductase (NM_080289) | 4 | T/U | -- | -- | 3' UTR |

*Analysis of the results of Ellis and colleagues (Ellis et al., 2005) reveal genes that are upregulated in a mouse with 2/3<sup>rd</sup> interstitial deletion of the long arm of the Y chromosome. UTRs of nine of these genes show homology to Y-transcripts. Most of the upregulated genes with homology within UTRs localize to the X-chromosome, with the exception of testatin and hydroxypyruvate reductase, which localize to autosomes. These results indicate putative interactions of the Y- chromosome with the X and the autosomes, via piRNAs. All the upregulated genes are expressed in testis (T), with few of them showing universal expression (U).*

**Figure 5.**
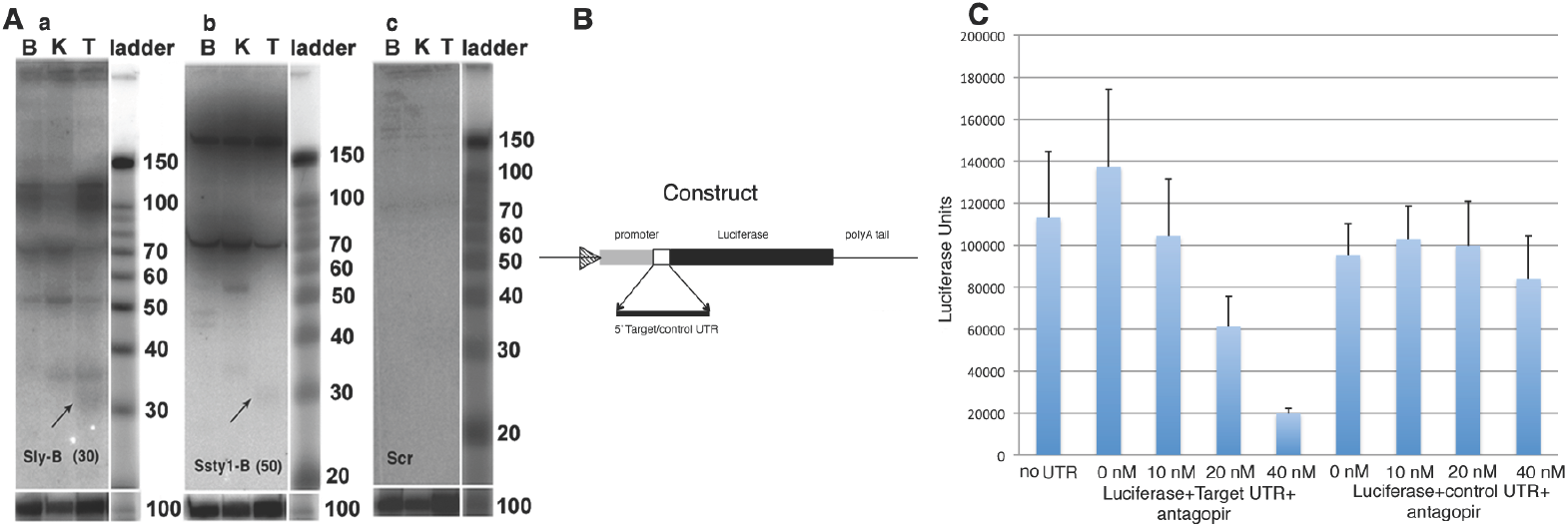
Genes regulated by Y chromosome have sequences homologous to piRNAs in their UTRs. **A**. **(a, b)** Shows hybridization with probes from Sly (Sly-B) and Ssty1 (Ssty1-B) that exhibit homology to UTRs of two upregulated genes (Ellis et al., 2005). Sly-B and Ssty1-B yielded ∼30 nt long testis-specific signals. **(c)** Scrambled miRNA probe served as the negative control. U6 was used as the loading control for all the blots. **B**. Cartoon representing Luciferase construct, where 5’UTR (RP11-130C19 gene) was inserted between promoter and coding region of Luciferase gene. **C**. Histogram representation of luciferase assay results from control and target UTRs containing homology to Y-transcripts with different concentrations of antagopirs. Figure shows decrease in Luciferase levels with increasing concentrations of antagopirs using the target UTR construct, whereas with the control UTR construct, antagopirs do not show a concentration dependent reduction in luciferase expression.

In a comparative sperm proteome analysis, using wild type XY^RIII^ strain of mice and a Y-deleted mutant, XY^RIII^qdel with a 2/3^rd^ interstitial deletion of the long arm of Y (3), we identified a few proteins that are differentially expressed between the two (19). Two of these proteins, calreticulin and superoxide dismutase (SOD) showed homology to Ssty2 in in their 3’ and 5’ UTRs respectively (Fig. 6)

**Figure 6.**
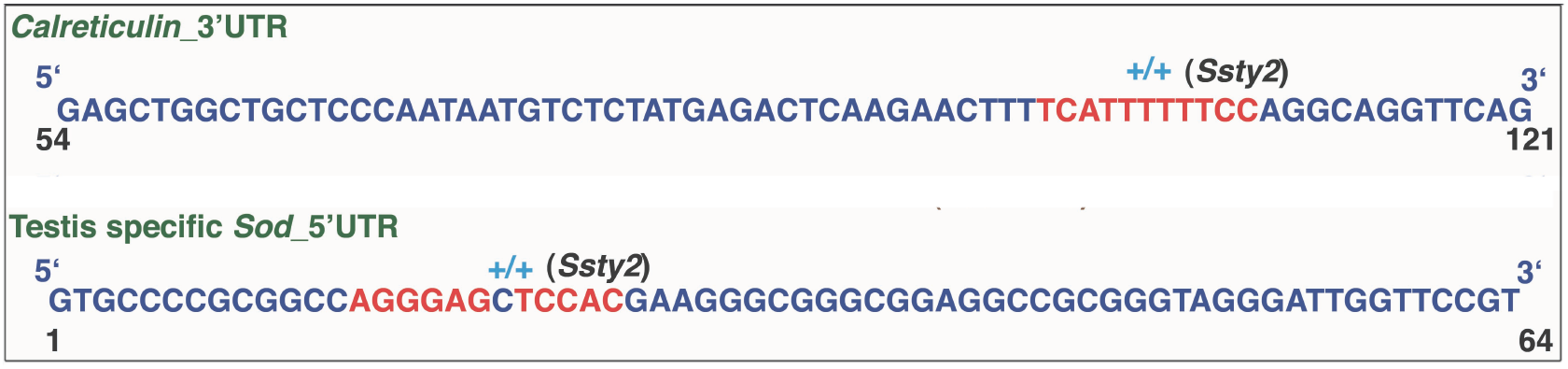
Homology to piRNAs in UTRs of deregulated genes. Two genes upregulated in XY^RIII^qdel mutant showed homology to Ssty2 in their UTRs

### Antagopirs downregulate reporter gene expression

Complementary oligonucleotides synthesized to piRNA sequences present in the 5’UTR of the gene localizing to clone RP11-130C19 was designated as antagopir. Increasing concentrations of antagopirs, i. e. 10 nM, 20 nM and 40 nM caused concentration dependent reduction in Luciferase expression (Fig. 5C). The antagopirs to 5’UTR of the gene from RP11-130C19 clone did not have an effect when the UTR from the *Gapdh* gene that lacks the piRNA homologous sequences was cloned 5’ to the Luciferase gene. Figure 5B is a schematic representation of 5’UTR reporter construct. Thus, the use of antagopirs to piRNA sequences present in UTR of the gene showed that piRNAs regulate gene expression.

## DISCUSSION

We establish here that small RNAs generated from the mouse Y muticopy gene transcripts are piRNAs based on (i) the identification of 30 nt RNAs mainly in testis, by Y-homologous sequences present in UTRs of different genes (ii) specific binding of these sequences to MIWI protein (iii) clustered organization on the mouse Y chromosome and (iv) differential expression from two DNA strands.

PIWI interacting RNAs (piRNAs) that range in size from 26-32 nt were reported initially from mouse testis by four different groups(17, 20-22). piRNAs are known to be generated from transposons and are involved in transposon silencing in genomes and germ cell maintenance(23-25). piRNA generating clusters have been localized to all the chromosomes except the Y in mammals(17, 22, 26, 27). The only report of generation of piRNAs from a Y chromosome is that of *Ste* (suppressor of stellate) locus in Drosophila(20, 24). Thus, with the present report of piRNAs from the multicopy gene transcripts from mouse Y, we elucidate a cluster of piRNAs on the mouse Y chromosome for the first time in mammals. piRNAs, which have roles in spine development and long-term memory have been reported from brain of mouse and Aplysia(28, 29). The expression of a Y chromosome-derived piRNA in mouse brain in the present study, is commensurate with reports of gene expression from the Y chromosome in mouse brain(30). Thus, we find that the mouse Y multicopy gene transcripts generate piRNAs in testis and brain.

The mouse Y multicopy gene transcripts that generate piRNAs do not contain secondary structures similar to that of precursors of miRNAs. In fact, previous investigators on using a few random piRNAs (along with their genomic flanks) and sequences from piRNA-generating loci did not identify secondary structures. Genomic sequences that were used as primary transcripts of piRNAs were not known(21). Watanabe and colleagues(22) predicted that ESTs localizing to the piRNA clusters could be precursors of piRNAs. Primary transcripts of piRNAs have been described from a piRNA generating loci in the silkworm BmN4 cell line using a GFP transgene(31). As we have identified piRNAs generated from the mouse Y transcripts, these are primary transcripts of piRNAs.

Deletion of mouse Y-heterochromatin results in upregulation of genes mainly from the X-chromosome(13). piRNAs from the Y chromosome transcripts have sequence homology to UTRs of genes that are upregulated in a Y-deletion mutant. Demonstration of piRNAs from a couple of these, suggests regulation of these genes by the Y-derived piRNAs. Modulation of Luciferase expression by the UTR of the gene from RP11-130C19 clone harboring piRNA homologous sequences further reiterates the regulatory role of these piRNAs. Reynard and Turner(32) hypothesized a gene on the male specific region of long arm of mouse Y (MSYq) that can regulate gene expression from the X chromosome. We propose that multicopy genes on the Y could regulate the expression from the X-chromosome via piRNAs.

Kim(33) hypothesized that, targets of piRNAs might be the PIWI complex and not RNA molecules per se. On the other hand, we propose that piRNAs from the Y chromosome may regulate a subset of genes in testis by view of homology to UTRs, analogous to digital controls. The functional significance of the two types of piRNAs, in +/+ and +/-orientation is however not clear. We find that both the sense and antisense strands of DNA produce piRNAs, although in unequal quantities. Many groups have observed differential expression of piRNAs from the two strands of DNA(17, 22, 33-35).

Based on the abundance of piRNAs in 3’UTRs of mRNAs, Robine and colleagues postulated the biogenesis of piRNAs from 3’UTRs(36). We find that approximately 37% of the piRNAs predicted from the Yq transcripts are complementary to the UTRs of different genes. Moreover, the genes that are upregulated in the Y-deletion mutant, have UTRs that are putative targets of the piRNAs encoded from the Y-repeat transcripts. This would suggest putative regulation of these genes by Y-chromosomal transcript derived piRNAs rather than biogenesis of piRNAs from their 3’UTRs. Further, microRNAs have been reported to regulate the expression of genes by binding to their 3’ UTRs, at regions of complementarity(37). MicroRNAs, are known to be generated from intergenic and intronic regions of genomes(38) regardless of homology to UTRs of different genes.

The protein-coding genes, degenerate pseudogenes, and repeat sequences on the mouse Y-heterochromatin are indispensable for spermatogenesis(3). The upregulation of gene expression from other chromosomes on deletion of the Y(13) suggests interaction between the Y chromosome and the X/autosomes. Our study provides evidence for the physical connection between mouse Y chromosome and transcripts from the rest of the genome via piRNAs. The multicopy *Ssty, Sly, Asty and Orly* genes localizing to the mouse Y chromosome, generate piRNAs that potentially regulate transcription and/or translation of Y-, X-chromosomal and autosomal genes besides coding for proteins.

This putative regulatory role of Y-chromosomal non-coding transcripts via piRNAs further strengthens the hypothesis of regulation of autosomal genes by noncoding transcripts from Y-heterochromatin repeats in testis(15), albeit by a different mechanism. A combinatorial approach of gene expression analyses using Y-deletion models together with characterization of noncoding RNAs from corresponding Y chromosomes may elucidate more examples of such interactions. The repeat sequences on Y-chromosomes appear to have been engineered for versatile regulatory roles.

## MATERIAL AND METHODS

RIII strain of mice(39) used was obtained as a gift from Prof. Paul S Burgoyne, and bred in our animal house facility; we have ethical committee clearance from our Institutional Review Board to use the mice (IAEC 65/28/2006).

### Small RNA Isolation

Total RNA was extracted from male (heart, spleen, kidney testis and brain) and female (ovary and brain) tissues of RIII mice using TRIZOL reagent (Invitrogen, Catalog Number: 15596-026). Total RNA was denatured at 65°C for 10 minutes, incubated with 10% PEG-8000 (Sigma-Aldrich, Catalog Number: P5413) and 5 M NaCl for 30 minutes on ice and centrifuged at 7,000 rpm for 7-10 minutes. The supernatant containing small RNA was precipitated overnight with 3 volumes of absolute alcohol and centrifuged at 13,000 rpm for 30 minutes. The small RNA pellet was washed with 80% ethanol and resuspended in RNase free water (Gibco-Life Technologies). The Small RNA was checked on 12% UREA PAGE (Urea, Biorad, Catalog Number: 161-0731) and quantitated using Nanodrop V-1000 (Thermo Fisher Scientific).

### Northern Blotting

20-50 µg of small RNA from each tissue was resolved on a 12% Urea PAGE gel in 0.5 x TBE running buffer and transferred onto Hybond N^+^ membrane. Decade marker (Ambion, Catalog Number: AM7778) was labeled and loaded according to the manufacturer’s instructions. 10-25 µM of each LNA-oligonucleotide probe (Exiqon), was end labeled for use as probes in hybridization buffer (5 x SSC, 5 x Denhardt’s and 1% SDS). Membranes were washed sequentially starting from room temperature to 70°C depending on the intensity of the signals. U6 was used as the loading control.

### Electrophoretic mobility shift assays

RNA oligonucleotides corresponding to Ssty (5’CCUCAUGAAGAAGAGGAGGAGGA3’), Sly (5’CAGUUAAAGAAAUGCAUGAGAA3’) and piR1 (5’UGACAUGAACACAGGUGCUCAGAUAGCUUU3’) were end labeled with γ-^32P^ATP and column purified using G-25 Sephadex (Sigma-Aldrich, Catalog Number: G25150) and quantitated on a scintillation counter. EMSA reactions were set up in a total volume of 25 μl using binding buffer (20 mM HEPES, 3 mM MgCl2, 40 mM KCl, 5% Glycerol, 2 mM DTT and RNase inhibitor (4U)), with MIWI protein (5 μg per reaction). MIWI was over-expressed from a recombinant construct in pAAV-IRES-hrGFP vector and purified using the FLAG tag. The purified MIWI protein was confirmed by Western blotting using MIWI (G82) antibody (Cell Signaling Technology, Catalog Number: 2079). Competitors i.e., unlabeled oligonucelotides (30 x concentration of hot oligo), MIWI antibody (90 ng) and Argonaute 3 antibody (100 ng) were added to the reaction, incubated for 1 h on ice, before addition of radio-labeled oligonucleotide (7000-10,000 cpm) and the entire mix was incubated on ice for another 30 min. EMSA was done on 5% native PAGE and image captured using FUJI phosphor Imager (FUJIFILM FLA-3000).

#### Luciferase assay

The 5’UTR sequences from the human clone RP11-130C19 and the *Gapdh* genes were amplified from mouse cDNA and cloned into the Luciferase vector. Either Luciferase gene alone or luciferase along with each UTR was cloned into pcDNA3.1 expression vector for assaying the effect of antagopirs (Figure 5B) on Luciferase reporter expression. Cotransfection experiments were done using the GC-1spg cell line (ATCC CRL-2053) and lipofectamine 2000 (Invitrogen) using protocols specified by the manufacturer. Cells were seeded in 48-well plates, 24 hours prior to transfection to obtain approximately 80% confluency. Each well was transfected with 50 ng of pcDNA3.1 plasmid containing either Luciferase gene alone or along with the cloned UTRs, 50 ng of the β-gal plasmid and varying concentrations of oligonucleotides complementary to the piRNA along with 0.5µl of lipofectamine 2000 in antibiotic and serum free DMEM (GIBCO). The complementary oligonucleotide to the piRNA has been designated as antagopir (5’ccucaugaagaagaggaggagga3’). The antagopirs were procured as RNA oligonucleotides from Eurofins Genomics India Pvt. Ltd, Bangalore, India. Different concentrations of antagopirs (0 nM, 10 nM, 20 nM and 40 nM/ well) were tested in the assay for their effect on the UTRs. Five hours-post transfection, the medium was replaced with complete growth medium. The cell extracts were prepared 24 hours-post transfection, using Reporter Lysis Buffer (Promega) and assayed for Luciferase activity in EnSpire 2300 multimode plate reader (Perkin Elmer). The Luciferase activity was normalized using β galactosidase. Three independent sets of experiments were done in triplicates.

## ACKNOWLEDGEMENTS

We wish to thank L. Singh for discussions, Jennifer Marshal Graves, S. Lakhotia, V. Radha and S. Tiwari for inputs during manuscript preparation, Lekha D Kumar for help with small RNA isolation, Arvind Kumar and Nitin Khandelwal for preparing the cell lysate for recombinant MIWI protein purification and Sunayana MR for help with editing the manuscript.

## AUTHOR CONTRIBUTIONS

RAJ – conceived the idea and directed the project, KM, AP – did the northern blots and other molecular biology experiments, AC, SK – did in silico analysis, NMP – did the gel shifts, HMR – initiated the experiments

## FUNDING

Department of Biotechnology, India [BT/PR 10707/AGR/36/596/2008] and Council of Scientific and Industrial Research (CSIR)’s intramural funding to RAJ.

## DISCLOSURE DECLARATION

The authors declare that they have no competing interests

---

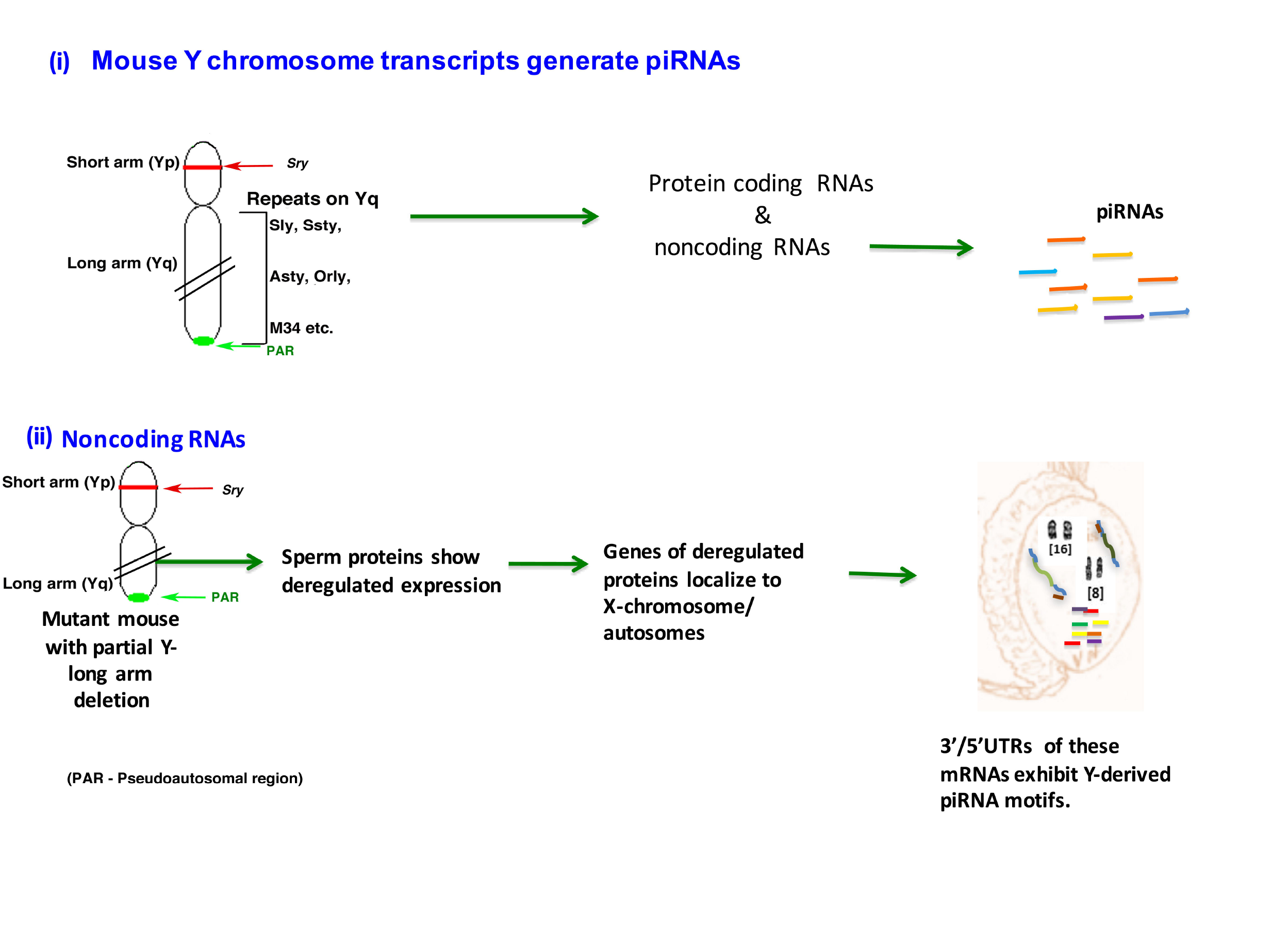

## REFERENCES

1. Kashimada K & Koopman P (2010) Sry: the master switch in mammalian sex determination. Development 137(23):3921–3930.

2. Burgoyne PS, Mahadevaiah SK, Sutcliffe MJ, & Palmer SJ (1992) Fertility in mice requires XY pairing and a Y-chromosomal “spermiogenesis” gene mapping to the long arm. Cell 71(3):391–398.

3. Toure A, et al. (2004) A new deletion of the mouse Y chromosome long arm associated with the loss of Ssty expression, abnormal sperm development and sterility. Genetics 166(2):901.

4. Nallaseth FS & Dewey MJ (1986) Moderately repeated mouse Y chromosomal sequence families present distinct types of organization and evolutionary change. Nucleic Acids Res 14(13):5295–5307.

5. Skaletsky H, et al. (2003) The male-specific region of the human Y chromosome is a mosaic of discrete sequence classes. Nature 423(6942):825.

6. Bonaccorsi S & Lohe A (1991) Fine mapping of satellite DNA sequences along the Y chromosome of Drosophila melanogaster: relationships between satellite sequences and fertility factors. Genetics 129(1):177–189.

7. Ellis PJ, Ferguson L, Clemente EJ, & Affara NA (2007) Bidirectional transcription of a novel chimeric gene mapping to mouse chromosome Yq. BMC Evol Biol 7:171.

8. Toure A, et al. (2004) A protein encoded by a member of the multicopy Ssty gene family located on the long arm of the mouse Y chromosome is expressed during sperm development. Genomics 83(1):140.

9. Cocquet J, et al. (2009) The multicopy gene Sly represses the sex chromosomes in the male mouse germline after meiosis. PLoS Biol 7(11):e1000244.

10. Toure A, et al. (2005) Identification of novel Y chromosome encoded transcripts by testis transcriptome analysis of mice with deletions of the Y chromosome long arm. Genome Biol 6(12):R102.

11. Soh YQ, et al. (2014) Sequencing the mouse Y chromosome reveals convergent gene acquisition and amplification on both sex chromosomes. Cell 159(4):800–813.

12. Ruiz-Orera J, Messeguer X, Subirana JA, & Alba MM (2014) Long non-coding RNAs as a source of new peptides. Elife 3:e03523.

13. Ellis PJ, et al. (2005) Deletions on mouse Yq lead to upregulation of multiple X- and Y-linked transcripts in spermatids. Hum Mol Genet 14(18):2705.

14. Riel JM, et al. (2013) Deficiency of the multi-copy mouse Y gene Sly causes sperm DNA damage and abnormal chromatin packaging. Journal of cell science 126(Pt 3):803-813.

15. Jehan Z, et al. (2007) Novel noncoding RNA from human Y distal heterochromatic block (Yq12) generates testis-specific chimeric CDC2L2. Genome Res 17(4):433–440.

16. Nishibu T, et al. (2012) Identification of MIWI-associated Poly(A) RNAs by immunoprecipitation with an anti-MIWI monoclonal antibody. Biosci Trends 6(5):248–261.

17. Girard A, Sachidanandam R, Hannon GJ, & Carmell MA (2006) A germline-specific class of small RNAs binds mammalian Piwi proteins. Nature 442(7099):199–202.

18. Reeder J, Steffen P, & Giegerich R (2007) pknotsRG: RNA pseudoknot folding including near-optimal structures and sliding windows. Nucleic Acids Res 35(Web Server issue):W320–324.

19. Hemakumar M. Reddy, 2,11 Rupa Bhattacharya,1,3,11 Zeenath Jehan,1,4 Kankadeb Mishra,1,5 Pranatharthi Annapurna,1,6 Shrish Tiwari,1 Nissankararao Mary Praveena,1 Jomini Liza Alex,1 Vishnu M Dhople,1,7 Lalji Singh,10 Mahadevan Sivaramakrishnan,1,8 Anurag Chaturvedi,1,9 Nandini Rangaraj1, Shiju Michael Thomas1, Badanapuram Sridevi,1 Sachin Kumar,1 Ram Reddy Dereddi1, Sunayana Rayabandla1, Rachel A. & Jesudasan1* (2018) Y chromosomal noncoding RNA regulates autosomal gene expression via piRNAs in mouse testis. BioRxiv.

20. Aravin A, et al. (2006) A novel class of small RNAs bind to MILI protein in mouse testes. Nature 442(7099):203–207.

21. Grivna ST, Beyret E, Wang Z, & Lin H (2006) A novel class of small RNAs in mouse spermatogenic cells. Genes Dev 20(13):1709–1714.

22. Watanabe T, et al. (2006) Identification and characterization of two novel classes of small RNAs in the mouse germline: retrotransposon-derived siRNAs in oocytes and germline small RNAs in testes. Genes Dev 20(13):1732–1743.

23. Klattenhoff C & Theurkauf W (2008) Biogenesis and germline functions of piRNAs. Development 135(1):3–9.

24. Siomi MC, Sato K, Pezic D, & Aravin AA (2011) PIWI-interacting small RNAs: the vanguard of genome defence. Nat Rev Mol Cell Biol 12(4):246–258.

25. Thomson T & Lin H (2009) The biogenesis and function of PIWI proteins and piRNAs: progress and prospect. Annu Rev Cell Dev Biol 25:355–376.

26. Aravin AA, Hannon GJ, & Brennecke J (2007) The Piwi-piRNA pathway provides an adaptive defense in the transposon arms race. Science 318(5851):761–764.

27. Grivna ST, Pyhtila B, & Lin H (2006) MIWI associates with translational machinery and PIWI-interacting RNAs (piRNAs) in regulating spermatogenesis. Proc Natl Acad Sci U S A 103(36):13415–13420.

28. Lee EJ, et al. (2011) Identification of piRNAs in the central nervous system. RNA 17(6):1090–1099.

29. Rajasethupathy P, et al. (2012) A role for neuronal piRNAs in the epigenetic control of memory-related synaptic plasticity. Cell 149(3):693–707.

30. Xu J, Burgoyne PS, & Arnold AP (2002) Sex differences in sex chromosome gene expression in mouse brain. Hum Mol Genet 11(12):1409.

31. Kawaoka S, et al. (2012) A role for transcription from a piRNA cluster in de novo piRNA production. RNA 18(2):265–273.

32. Reynard LN & Turner JM (2009) Increased sex chromosome expression and epigenetic abnormalities in spermatids from male mice with Y chromosome deletions. J Cell Sci 122(Pt 22):4239–4248.

33. Kim VN (2006) Small RNAs just got bigger: Piwi-interacting RNAs (piRNAs) in mammalian testes. Genes Dev 20(15):1993–1997.

34. Houwing S, et al. (2007) A role for Piwi and piRNAs in germ cell maintenance and transposon silencing in Zebrafish. Cell 129(1):69–82.

35. Yamamoto Y, et al. (2013) Targeted gene silencing in mouse germ cells by insertion of a homologous DNA into a piRNA generating locus. Genome Res 23(2):292–299.

36. Robine N, et al. (2009) A broadly conserved pathway generates 3’UTR-directed primary piRNAs. Curr Biol 19(24):2066–2076.

37. Barrett LW, Fletcher S, & Wilton SD (2012) Regulation of eukaryotic gene expression by the untranslated gene regions and other non-coding elements. Cell Mol Life Sci 69(21):3613–3634.

38. Kim VN & Nam JW (2006) Genomics of microRNA. Trends Genet 22(3):165–173.

39. Conway SJ, et al. (1994) Y353/B: a candidate multiple-copy spermiogenesis gene on the mouse Y chromosome. Mamm Genome 5(4):203.

